# Genetic dissection of DNA damage response pathways reveals divergent modulation of PD-1 blockade response in colorectal cancer

**DOI:** 10.64898/2026.10.03.756428

**Authors:** Ferran Fece de la Cruz, Edmond Wong, Haley Barnes, Katerina Fella, Jacquelyn Curtis, Sarah Phat, David T Myers, Parasvi S Patel, Jonathan Braverman, Omer H Yilmaz, Ryan B Corcoran

## Abstract

Although MSI-H/dMMR colorectal cancers and the rarer subset with pathogenic *POLE* exonuclease domain mutations can benefit from immune checkpoint blockade, MSS colorectal cancer remains largely refractory, raising the question of whether other DNA damage response defects may also confer sensitivity to immunotherapy. Here, we performed a systematic *in vivo* screen of recurrently altered DNA damage response (DDR) genes to determine whether distinct repair defects modulate response to PD-1 blockade in MSS colorectal cancer. Using a genome-edited murine model in immunocompetent mice, we found that DDR perturbations produced heterogeneous outcomes, indicating that the immunologic consequences of DNA repair deficiency may be pathway-specific. Loss of *Brca1*, *Brca2*, *Atm*, and *Fanca* enhanced response to anti–PD-1 therapy, whereas *Atr* and *Msh6* deficiency were associated with poor benefit. Brca1/2-deficient tumors showed the strongest regressions, supporting a link between core homologous recombination deficiency and increased immune sensitivity. In an independent cohort of colorectal cancer patients treated with ICB from the MSK-IMPACT Clinical Sequencing Cohort, *BRCA1* alterations were associated with improved survival status, while *FANCA* alterations showed a similar trend. Because ATM is therapeutically targetable, we evaluated pharmacologic ATM inhibition in MSS colorectal cancer. Sustained ATM inhibition induced immune-related transcriptional programs *in vitro* and, when administered as a pre-treatment *in vivo*, enhanced the antitumor activity of PD-1 blockade. Together, these findings suggest that selected DDR alterations may represent candidate biomarkers of immunotherapy sensitivity, while ATM inhibition may provide a rational combination strategy to enhance PD-1 blockade in MSS colorectal cancer.

## Introduction

Immune checkpoint blockade (ICB) therapies have transformed the treatment landscape of multiple malignancies, providing durable clinical responses in subsets of patients with melanoma, non–small cell lung cancer, and other solid tumors(1, 2). However, in colorectal cancer (CRC), the clinical success of programmed cell death-1 (PD-1) blockade has been largely confined to tumors harboring microsatellite instability–high (MSI-H) or mismatch repair–deficient (dMMR) phenotypes(3, 4). These tumors exhibit high tumor mutational burden (TMB) and elevated neoantigen loads, fostering immune infiltration and sensitivity to PD-1 inhibition(5). In contrast, the majority of CRCs are microsatellite stable (MSS) and mismatch repair–proficient (pMMR), characterized by limited immune infiltration and intrinsic resistance to ICB(6). The molecular and genetic determinants underlying this immunologic divergence remain incompletely understood, owing in part to the scarcity of physiologically relevant preclinical models that accurately reflect the genetic and immunologic complexity of human CRC.

Current *in vivo* studies exploring immunotherapy response in CRC predominantly employ two murine models: CT-26 and MC-38. Both were derived from tumors induced in mice exposed to colitis- or carcinogen-promoting agents such as N-nitroso-N-methylurethane (NMU) or dimethylhydrazine(7–9). While these lines have been instrumental in defining general immunologic principles, they incompletely represent the genomic architecture and evolutionary path of human colorectal carcinogenesis. CT-26 lacks canonical *Apc* or *Tp53* alterations, and its molecular and immunologic features do not fully recapitulate human pMMR colorectal cancer, whereas MC-38 more closely resembles hypermutated MSI-H disease(10, 11). Consequently, these models do not capture the sequential accumulation of oncogenic lesions—from *APC* loss to *KRAS* activation, *TP53* inactivation, and *SMAD4* disruption—that characterize the adenoma–carcinoma transition(12). The limited genetic fidelity of these models, coupled with their variable immunogenicity, constrains mechanistic exploration of tumor–immune crosstalk and hinders development of rational combination therapies.

To address these limitations, we developed a genetically defined murine colorectal cancer model, termed AKPS (**Fig. 1A**), that integrates the hallmark driver mutations observed in human CRC: *Apc* knockout (KO), oncogenic *Kras*^G12D^, *Tp53* KO, and *Smad4* KO. Together, these alterations recapitulate the core signaling disruptions governing colorectal epithelial transformation, including deregulation of Wnt, Ras/Mapk, Tp53, and Tgfβ pathways(13). Importantly, AKPS cells maintain robust tumorigenicity in immunocompetent hosts and can be efficiently engineered to express Cas9 and single guide RNAs (sgRNAs) without compromising their proliferative or tumor-forming capacity(14). This genetic tractability positions the AKPS model as a versatile and physiologically relevant platform for functional interrogation of tumor-intrinsic mechanisms shaping immune response in CRC.

**Figure 1.**
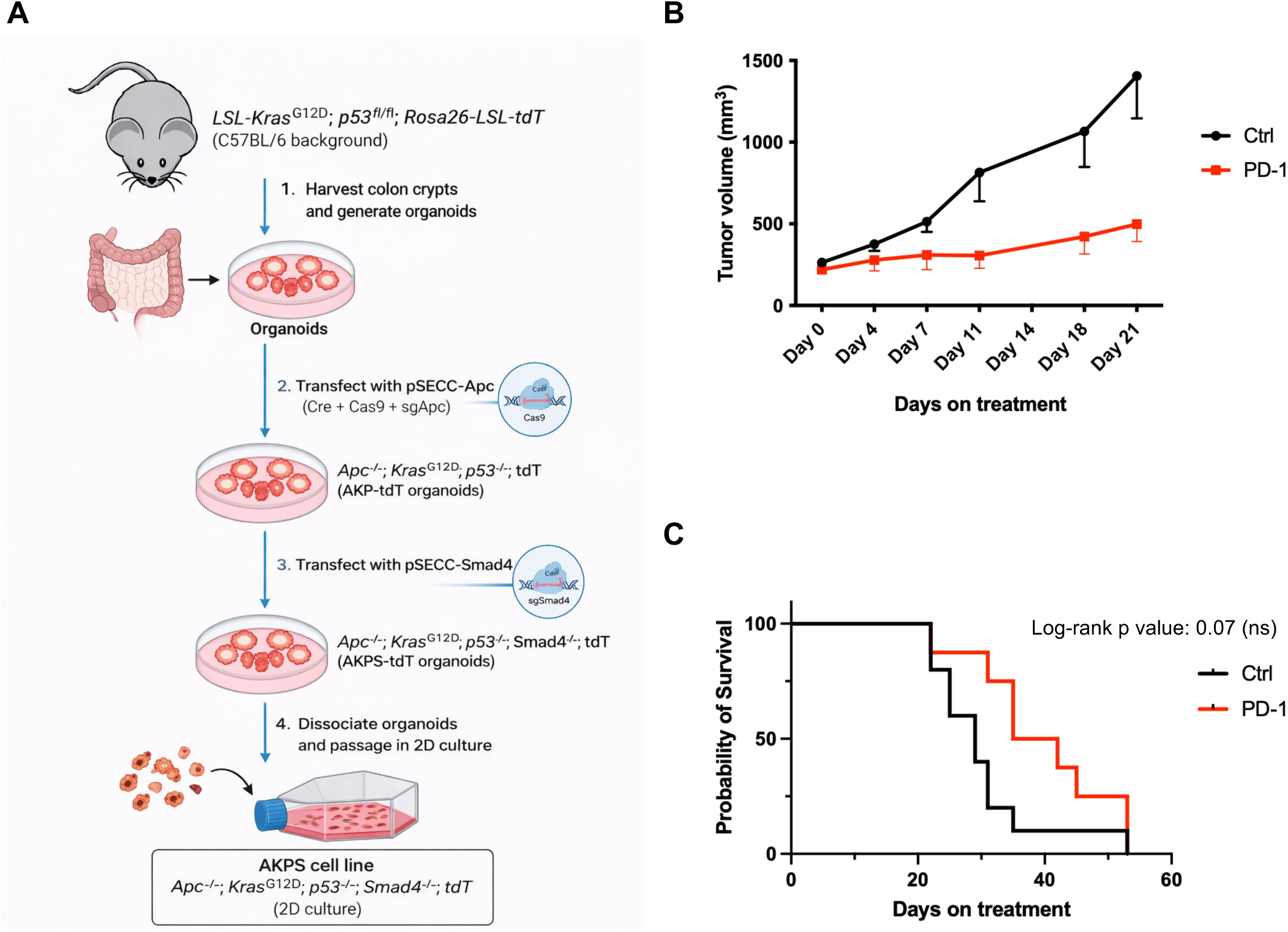
Generation of AKPS cells and characterization *in vivo*. (**A**) Flowchart describing how AKPS cells were established from mouse colon crypts. (**B**) Growth of AKPS cells implanted in the flanks of C57BL/6 mice and treated with anti-PD-1 antibodies (10 mg kg^−1^ twice per week) or an IgG control. Shown is the mean and standard error (n=8, Mann-Whitney test p=0.09, ns). (**C**) Kaplan-Meier survival curve of mice in **B**. IgG control vs PD-1 p=0.07, Log rank Mantel-Cox test.

Genomic instability is increasingly recognized as a double-edged regulator of tumor immunogenicity(15). Defects in the DNA damage response (DDR) machinery can potentiate immune recognition by increasing neoantigen generation, activating cytosolic DNA sensing pathways, and inducing type I interferon signaling(16, 17). Conversely, certain DDR alterations may promote immune evasion by impairing antigen presentation, perturbing interferon signaling, or fostering a chronically inflamed yet immunosuppressive tumor microenvironment(18). Clinically, dMMR tumors exhibit striking responses to PD-1 blockade, whereas alterations in non-MMR DNA repair pathways, particularly HRR, have shown context-dependent associations with immunogenicity(19, 20). Despite these observations, a comprehensive understanding of how specific DDR pathways influence PD-1 blockade response in CRC remains elusive.

## Materials and methods

### AKPS cell line generation

Colon crypts were harvested from LSL-Kras^G12D^; p53 fl/fl; Rosa26-LSL-tdT mice on a C57BL/6 background and organoids were generated. Organoids were transfected with a pSECC-Apc plasmid (carrying Cre, Cas9 and a sgRNA targeting Apc) to generate Apc^-/-^; Kras^G12D^; p53^-/-^; tdT organoids (AKP-tdT). AKP-tdT organoids were selected by Wnt withdrawal to select for Apc^-/-^ and addition of Nutlin-3 to select for p53^-/-^. Cells were sorted by flow cytometry based on a >95% tdT expression. Selected AKP-tdT organoids were transfected with pSECC-Smad4 and selected for Smad4^-/-^(AKPS-tdT) by passaging in TGFb-supplemented medium. AKPS-tdT organoids were dissociated and passaged as a 2D cell culture.

### Cell culture and general reagents

AKPS cells were cultured in Advanced Dulbecco’s Modified Eagle Medium (DMEM)/F12 media supplemented with 10% fetal bovine serum (FBS) and 2mM GlutaMAX (Thermo Fisher Scientific). HEK293T cells obtained from ATCC were maintained in DMEM with 10% FBS. All cells were grown in the presence of penicillin–streptomycin at 37°C and 5% CO_2_. Mycoplasma testing was performed routinely using a Mycoplasma PCR detection kit (ABM, #G238). All cell lines used tested negative for *Mycoplasma*.

Lentiviral particles were produced in HEK293T cells. LentiCas9-blast (#52962) and LentiGuide-puro (#52963) vectors were obtained from Addgene. Cas9 was stably integrated into AKPS cells by lentiviral transduction followed by blasticidin selection. sgRNAs targeting each gene of interest was stably integrated into AKPS Cas9-expressing cells by lentiviral transduction followed by puromycin selection. Anti-PD-1 antibody for animal studies was kindly provided by Gordon Freeman.

### Dose-response experiments

Viability assays were carried out using the CellTiter-Glo assay (Promega). Cells were counted and seeded in 96-well plates in triplicates. The next day, compounds were added, and 3 to 4 days later, cell viability was assessed.

### Western Blot analysis

Lysates were prepared as described previously(21). Mouse BRCA1 antibody was kindly provided by Andre Nussenzweig (NCI, Bethesda, USA). Mouse BRCA2 antibody was kindly provided by Hyunsook Lee (Seoul National University, Seoul, South Korea). Mouse PALB2 antibody was kindly provided by Bing Xia (Rutgers Cancer Institute, New Jersey, USA). CDK12 antibody (#ABE1861) was ordered from Millipore Sigma. The following antibodies were purchased from Cell Signaling Technology: ATM (#2873S), ATR (#13934S), ARID1A (#12354S), MSH2 (#2017S), MSH6 (#3995S), Tubulin (#3873S) and phospho-Chk2 (#2661). MLH1 antibody (#ab92312) and B2M antibody (#ab75853) were ordered from Abcam. FANCA antibody (#PA5-105723) was obtained from ThermoFisher.

### Quantitative real-time PCR

RNA was extracted using a RNeasy kit (Qiagen) per the manufacture’s protocol. Reverse transcription was performed using qScript cDNA SuperMix (Quantabio). Quantitative real-time PCR was performed employing the FastStart Universal SYBR Green Master (Rox). Analysis was carried out in triplicates, using *GAPDH* as a control gene.

### Generation of isogenic cell lines

A single cell Cas9-expression clone was isolated and transduced with sgRNAs targeting each gene of interest followed by puromycin selection (Sequences for all sgRNA can be found in **Suppl Table 1**). Isogenic cell lines were passaged *in vitro* for 4-5 weeks before being injected into immunocompetent mice. All of them were validated using western blotting.

### Whole-exome sequencing analysis

Whole-exome sequencing was performed on AKPS-derived knockout cell populations and an empty-vector control AKPS line after 4–5 weeks of *in vitro* expansion. Paired-end clean reads were aligned to the mouse reference genome mm10 using Burrows-Wheeler Aligner (BWA)(22). Alignment files were generated in BAM format, sorted with SAMtools(23), and processed with Picard to merge BAM files from the same sample and mark duplicate reads. Coverage and sequencing depth were calculated from the final processed BAM files.

Reference-relative SNPs and indels were detected and filtered using GATK(24). Somatic variants were identified using the empty-vector control AKPS line as the matched reference. Somatic SNVs were called with MuTect(25), and somatic indels were called with Strelka(26). Somatic copy-number variants were detected using Control-FREEC(27). Variant annotation was performed using ANNOVAR(28).

### Animal studies

AKPS cells were resuspended in Advanced DMEM/F12 medium (without FBS) and Corning Matrigel in a 1:1 ratio and then injected (7 × 10^5^ cells per injection) into the flanks of 12-week-old male C57BL/6 mice (Charles River Laboratories). When the tumor size reached ∼200 mm^3^, mice were randomized into anti-PD-1 (10 mg kg^−1^ twice per week, intraperitoneal injection) or immunoglobulin G isotype control (BioXcell, intraperitoneal injection). Tumors were measured using calipers twice per week and tumor volume was estimated using *V*=1/2(*L* × *W*^2^), where *L* is the longest dimension (length) and *W* is width (shortest dimension). Animal studies and procedures were performed in accordance with the institutional guidelines of Massachusetts General Hospital, and all experiments were conducted according to institutional animal care and use committee-approved protocols.

## Results

### Characterization of AKPS cell model *in vivo*

To determine whether AKPS represents a suitable model for studying response to immune checkpoint blockade in colorectal cancer, we first assessed its tumor growth kinetics and response to anti–PD-1 therapy (**Fig. 1**). As expected in a context of MSS/pMMR colorectal cancer, while there was some initial therapeutic benefit in delaying tumor growth, the PD-1-treated animal cohort did not experience a significant survival benefit compared to the control animal group (**Fig. 1B-C**). Additionally, we rarely observed tumor regressions in this model in response to immune checkpoint blockade, recapitulating the current clinical observations in colorectal cancer patients.

One limitation when studying responses to immunotherapy using genetically engineered cancer cell models is that the expression of Cas9 or other CRISPR components can itself elicit immune recognition in immunocompetent mouse models. As Cas9 is derived from bacterial proteins, its expression in tumor cells may generate neoantigenic peptides that are presented by major histocompatibility complex (MHC) molecules, leading to T-cell–mediated immune responses and potential tumor rejection independent of the engineered genetic alterations. To address this limitation, we generated several Cas9-expressing AKPS clones (**Suppl. Fig. 1A**) and evaluated their tumorigenic potential and response to PD-1 blockade in immunocompetent mice. All AKPS-Cas9 clones tested could form tumors *in vivo*, although some variability in tumor growth kinetics was observed across clones. Among these, AKPS-Cas9 clone 2 showed robust engraftment *in vivo* and, relative to the other clones tested, provided the most favorable balance between reliable tumor growth and efficient genome editing (**Suppl. Fig. 1B**). Based on these properties, we selected AKPS-Cas9 clone 2 for subsequent genetic manipulation and functional studies. Together, the AKPS model recapitulates a well-defined genetic model of colorectal cancer, displays limited-in-time response to PD-1 blockade and is amendable to genetic perturbations to mimic the complex heterogeneity of colorectal cancer.

### An *in vivo* screen of genetically defined DNA repair defects reveals divergent responses to PD-1 blockade

Using the genetically tractable AKPS system, we systematically investigated how individual DNA damage response (DDR) pathways contribute to antitumor immune modulation and influence response to immune checkpoint blockade. We generated an isogenic panel of AKPS-derived knockout cell lines targeting the most frequent altered DDR genes in colorectal cancer: *ARID1A* (8.2%), *ATM* (7.4%), *BRCA2* (5.9%), *ATR* (3.6%), *MSH6* (3.3%), *FANCA* (3%), *CDK12* (2.9%), *BRCA1* (2.8%), *MSH2* (2.5%) and *PALB2* (2.1%) (**Suppl. Fig. 2**). Another frequently mutated gene *SETD2* (4.4%) was excluded due to spontaneous regressions in some SETD2-KO tumors. Our selection for this *in vivo* screen included genes involved in mismatch repair (MMR), homologous recombination-mediated repair (HRR), cell cycle checkpoints, and DNA damage signaling, thereby capturing a spectrum of alterations that can shape tumor mutational evolution. Although loss of these genes does not necessarily compromise cell viability, cells deficient in these pathways are expected to accumulate additional genomic aberrations over time as a consequence of impaired DNA repair.

AKPS engraftment in immunocompetent mice achieves a nearly 100% efficiency and tumors reach the adequate size to start treatment within two weeks post-implantation. We reasoned that injecting the engineered cells right after inactivating a particular gene would not be sufficient for each isogenic cell line to accumulate a distinct evolutionary trajectory mediated by the inactivation of a particular DDR pathway. Therefore, these isogenic populations were expanded *in vitro* for 4-5 weeks to allow them to accumulate additional genomic alterations driven by their respective repair deficiencies (**Fig. 2A**). AKPS cell division time is slightly under 24 hours, so we estimated that around 40 cell divisions occurred after the gene of interest was disrupted and we injected the cells to the flanks of immunocompetent animals, allowing our isogenic cell panel to achieve differentiated trajectories in terms of accumulated DNA damage.

**Figure 2.**
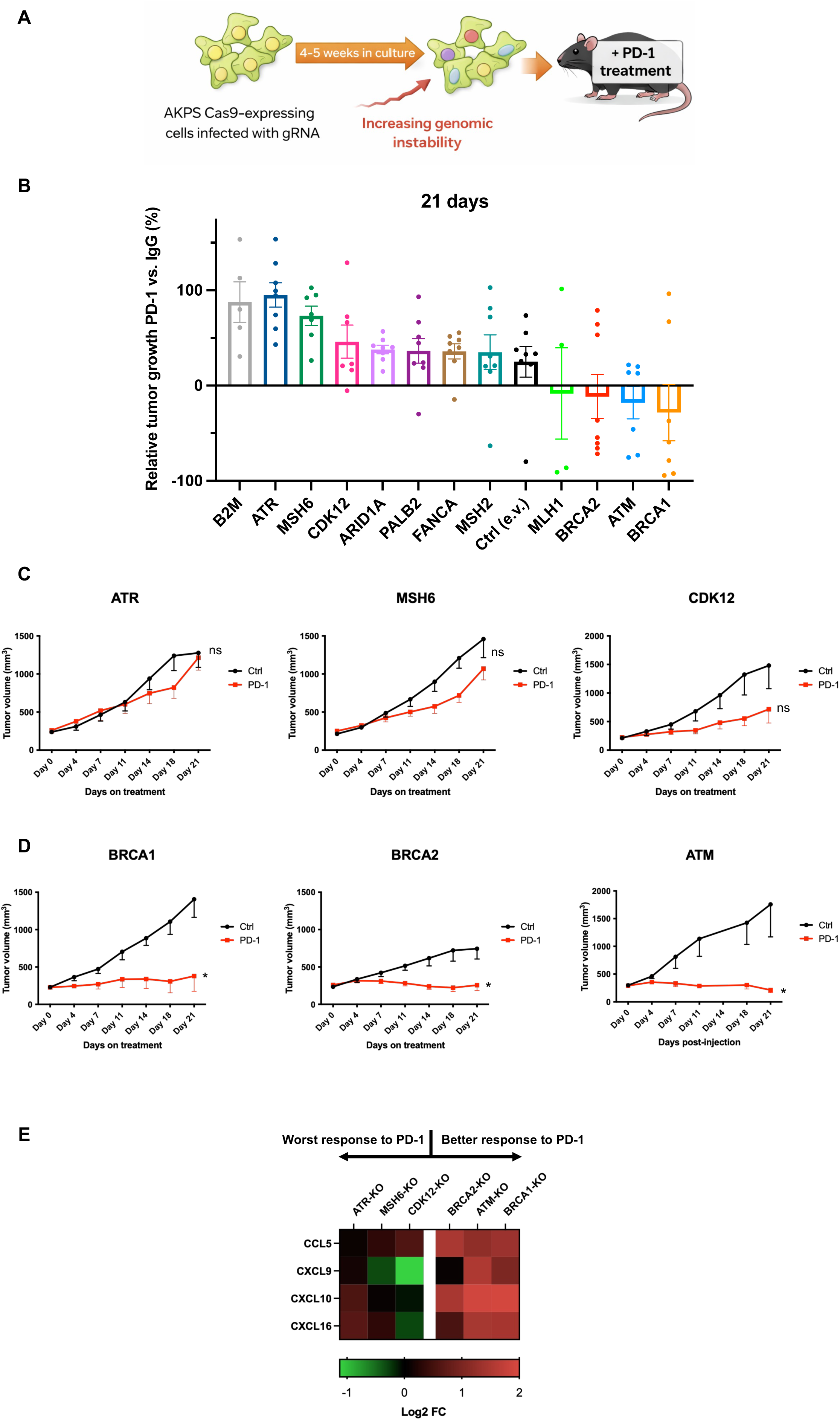
An *in vivo* screen reveals divergent responses to PD-1 blockade among tumors with distinct DDR deficiencies. (**A**) Diagram depicting how isogenic cell lines were generated in AKPS-Cas9 expressing cells before being injected into immunocompetent mice. (**B**) Relative tumor growth in PD-1 treated animals compared to animals treated with IgG control at day 21 of treatment. Shown is the mean and standard error, each dot represents an independent animal. (**C**) Growth of AKPS tumors infected with gRNAs targeting *Atr*, *Msh6* or *Cdk12*, respectively. Shown is mean and standard error (n=7-8). (**D**) Growth of AKPS tumors infected with gRNAs targeting *Brca1*, *Brca2* or *Atm*, respectively. Shown is mean and standard error (n=7-8). * p<0.05, Mann-Whitney test. (**E**) Heatmap visualizing gene expression changes in the indicated AKPS-KO isogenic cell lines. Data are normalized to control empty vector AKPS cells (n=3).

To confirm that the edited populations had acquired measurable genomic alterations during this expansion period, we characterized the AKPS-DDR panel by whole-exome sequencing (WES) prior to implantation. Using an empty-vector control AKPS line as the reference for somatic variant calling, we detected genotype-specific accumulation of small variants and copy-number alterations across the panel (**Suppl. Fig. 3**). Total somatic small-variant burden varied across knockout populations, with differences in the relative contribution of SNPs and indels (**Suppl. Fig. 3A, C, and D**). WES-normalized protein-altering coding mutational burden was also variable across genotypes, indicating that the evolved populations differed not only in total variant number but also in the fraction of alterations predicted to affect coding sequence (**Suppl. Fig. 3B**). In parallel, copy-number analysis revealed distinct CNV architectures, including differences in total CNV event number and the genomic territory affected by copy-number gains or losses (**Suppl. Fig. 3E**). These data confirmed that a relative short-term *in vitro* evolution after DDR gene disruption was sufficient to generate measurable, genotype-associated genomic divergence within the isogenic AKPS panel, providing a rationale to test whether these distinct evolutionary states influenced a response to PD-1 blockade *in vivo*.

Our AKPS-DDR panel of isogenic cells engrafted to similar extend as the AKPS parental cell line, except for *Brca2*-KO cells which tended to grow tumors at a slower pace (**Fig. 2C-D**). Nonetheless, all cell lines achieved adequate tumor size allowing treatment with anti-PD-1 antibodies. In contrast to the initial limited-in-time favorable response observed in parental AKPS, some of our DDR-deficient cell lines presented with a differential response to PD-1 blockade, observing both marked resistance as well as tumor regressions at day 21 of treatment (**Fig. 2B and Suppl. Fig. 4**). AKPS tumors with deficiency in either *Atr* or *Msh6* accounted for the less responsive ones (**Fig. 2C**), with a growth pattern comparable to our positive control for resistance to anti-PD-1 therapy (*B2m*-KO) (**Fig. 2B**). It has been previously described that ATR loss-of-function mutations in melanoma harness the immune system to accelerate tumor growth(29). On the other hand, ATR inhibition has been associated to a better outcome to checkpoint immunotherapy in lung cancer and melanoma(30, 31). Our panel of isogenic AKPS-KO cell lines is expected to contain a heterogeneous pool of edited clones with distinct fitness to proliferate both *in vitro* and *in vivo*. Therefore, we can’t exclude the possibility that we selected for clones containing a similar pattern of loss-of-function mutations leading to resistance.

Among the best responders to anti-PD1 therapy, we observed an agreement in both cell lines depleted for *Brca1* and *Brca2*, suggesting a role for HRR-deficiency inducing a most immune responsive microenvironment. We also observed a favorable profile of response in ATM-KO tumors, even if they were among the most aggressive tumors in our isogenic panel (**Fig. 2D**). Remarkably, tumor regressions were observed in these three groups (5/7 for *Brca1*-KO, 5/8 for *Brca2*-KO and 3/8 for *Atm*-KO). The divergence in response between *Atr*-KO and *Atm*-KO tumors was noteworthy, especially since these two proteins are both involved in response to DNA strand breaks and loss-of-function of one can be to some extent compensated it by the other.

Tumor-intrinsic factors can reshape the tumor microenvironment by attracting or preventing immune cell infiltration. In this regard, we hypothesized that transcriptional changes derived from different patterns in genome instability could determine the differential response to PD-1 blockade observed in our panel of isogenic cell lines. Indeed, the three more sensitive cell lines (*Brca1*-KO, *Brca2*-KO and *Atm*-KO) presented upregulation of immune cell attractant chemokines such as CCL5, CXCL9, CXCL10 and CXCL16 (**Fig. 2E**). In contrast, we observed a downregulation of these genes in the group of cell lines that showed less benefit to anti-PD-1 treatment (*Atr*-KO, *Msh6*-KO and *Cdk12*-KO).

Together, these divergent outcomes suggest that DNA repair deficiency may not define a uniform immunologic state. Instead, the specific DDR pathway disrupted may shape whether tumors evolve toward increased sensitivity or resistance to PD-1 blockade. (**Suppl. Fig. 5**).

### Biomarkers of response to PD-1 blockade in colorectal cancer

The results of the *in vivo* screen demonstrated that tumors lacking *Brca1* or *Brca2* exhibit a remarkable response to PD-1 blockade (**Fig. 2B**), consistent with the profound homologous recombination (HR) deficiency and accumulation of genomic aberrations caused by loss of these core HR repair factors. In contrast, disruption of other HR pathway regulators, such as *Palb2*, produced substantially weaker effects, with only one tumor regression observed at day 21 (**Fig. 2B**). Tumors harboring *Palb2* loss showed responses more comparable to those observed in wild-type control tumors, suggesting that this perturbation induces a less pronounced HR deficiency–associated genetic damage state. Together, these findings indicate that loss of the central *Brca1/Brca2* HR machinery generates a stronger immunogenic phenotype than perturbation of upstream or accessory HR regulators.

Interestingly, animals injected with *Brca1*-KO cells achieved a 71% success on tumor regression with a notable durable response up to 60 days, when the PD-1 treatment was discontinued. Out of the five responders, three presented with a complete response without tumor recurrence beyond 90 days after treatment was initiated (**Fig. 3A**). As previously mentioned, tumor regressions in response to immune checkpoint blockade are rare in the AKPS model, indicating the strong immunogenic effect on colorectal cancer cells evolving under BRCA1 deficiency.

**Figure 3.**
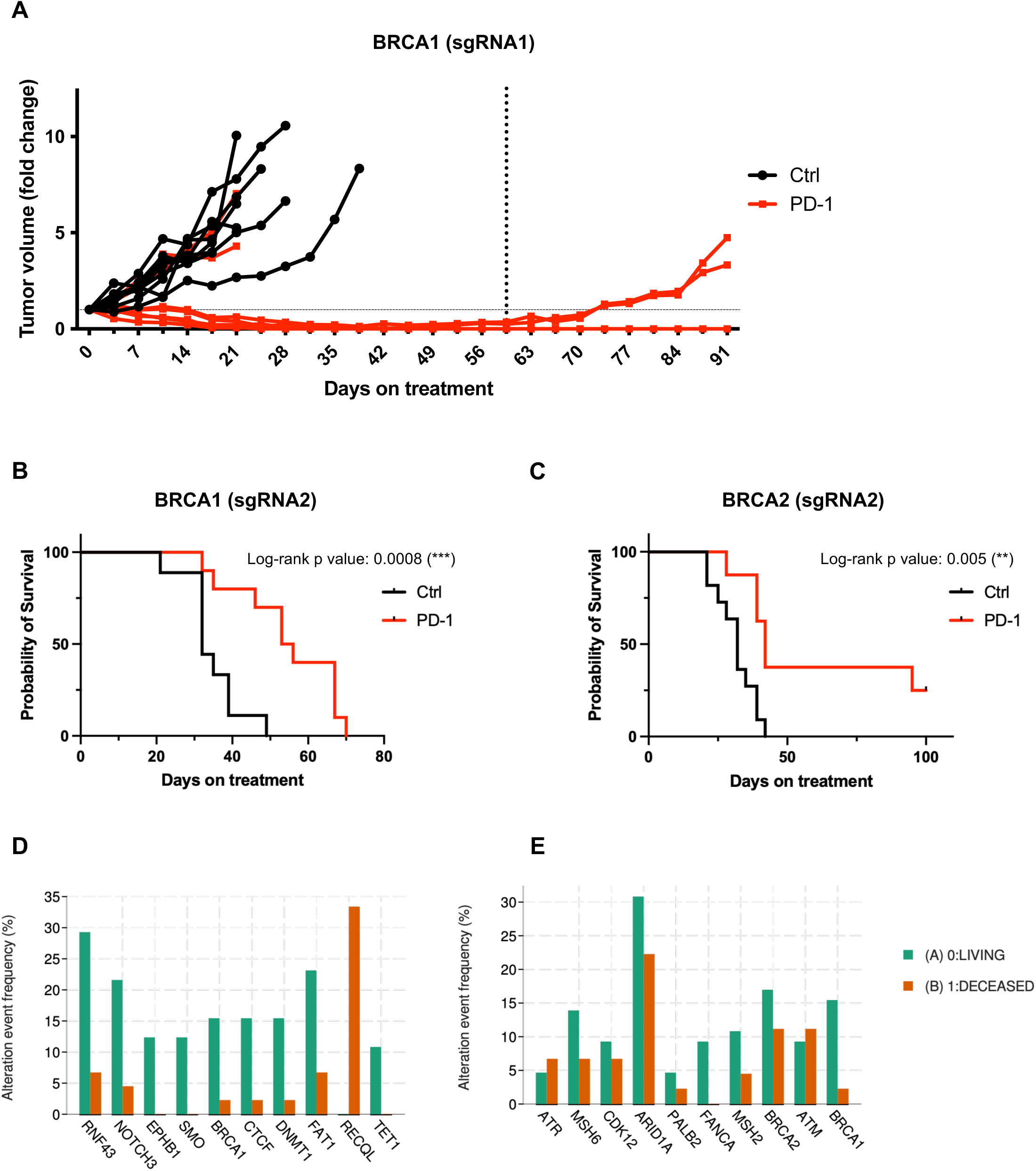
BRCA1/2 deficiency represents a potential biomarker of response to PD-1 blockade in colorectal cancer. (**A**) Fold change in tumor growth of AKPS *Brca1*-KO cells treated with anti-PD-1 antibodies (10 mg kg^−1^ twice per week) or an IgG control. Every line represents an independent animal (n=7-8). Treatment was withdrawn at day 60 (dotted vertical line). (**B**) Kaplan-Meier survival curve of mice injected with *Brca1*-sgRNA2 cells and treated with anti-PD-1 antibodies (10 mg kg^−1^ twice per week) or an IgG control (n=9-10). p<0.005, Log rank Mantel-Cox test. (**C**) Kaplan-Meier survival curve of mice injected with *Brca2*-sgRNA2 cells and treated with anti-PD-1 antibodies (10 mg kg^−1^ twice per week) or an IgG control (n=9-11). p=0.005, Log rank Mantel-Cox test. **(D–E)** Matched genomic alteration and survival status data were obtained for 110 colorectal cancer patients treated with immune checkpoint blockade from the MSK-IMPACT cohort accessed through cBioPortal. Altered gene status is defined as the presence of a mutation, structural variant, or gene fusion in the indicated gene. Panels show the 10 genes most significantly associated with survival status in this cohort (**D**) and the 10 genes included in our *in vivo* screen (**E**).

In a subsequent independent validation experiment using alternative sgRNAs to target either *Brca1* or *Brca2* (**Suppl. Fig. 6**), both genotypes again showed improved overall survival, confirming the association between homologous recombination deficiency and enhanced therapeutic response (**Fig. 3B-C**). Notably, whereas the initial experiment showed a stronger effect for *Brca1* loss, the validation cohort showed a more pronounced benefit in animals bearing *Brca2*-deficient tumors, with a greater fraction of these mice achieving prolonged survival. These differences likely reflect sgRNA-dependent variation in editing efficiency or allelic disruption, but they may also be consistent with the distinct functions of BRCA1 and BRCA2 within homologous recombination repair. Specifically, BRCA1 functions at early stages of homologous recombination, including DNA end resection and repair pathway choice, whereas BRCA2 acts more directly in RAD51 loading and strand invasion. These nonredundant roles may result in differences in the extent or persistence of repair-associated genomic alterations following gene loss.

The MSK-IMPACT database contains clinical data and targeted sequencing results from 1,662 patients treated with immune checkpoint blockade, including 110 colorectal cancer patients, with a median follow-up of 19 months (range 0-80) from the date of first ICI treatment(32). Tumor mutational burden (TMB) and overall survival analysis for the colorectal cancer cohort can be seen in **Suppl. Fig 7A**. The majority of colorectal cancer patients was treated with anti-PD-1 regime as monotherapy (90%) or in combination with an anti-CTL4 regime (9.1%), therefore we investigated whether tumors with alterations (mutations, structural variations or fusions) in our gene panel could be matched to response to PD-1. Among the top ten altered genes most significantly associated with survival status, *BRCA1* alterations were more frequent in patients who were alive than in those who were deceased (**Fig. 3D**). Higher TMB has been associated with improved outcomes in patients receiving immunotherapy. Among the 110 colorectal cancer patients analyzed, approximately one-third had high TMB, defined as >24 mutations/Mb (**Suppl. Fig. 7A**). Notably, all 10 patients harboring *BRCA1* alterations exhibited high TMB, with values ranging from 38.62 to 178.44 mutations/Mb (**Suppl. Fig. 7B**). Interestingly, 6 of these 10 patients did not harbor alterations in canonical hypermutation-associated genes, including *MLH1* or the *POLE* exonuclease domain. (**Suppl. Table 2**). These observations suggest that *BRCA1* alterations may be enriched among tumors with elevated TMB, although additional analyses will be required to determine whether *BRCA1* loss directly contributes to this phenotype or co-occurs with other processes driving mutational accumulation.

Overall, within the genes included in our panel, only *BRCA1* alterations were significantly associated with improved survival following immune checkpoint blockade (ICB) in the MSK-IMPACT cohort (**Fig. 3E**). *BRCA1* alterations were identified in 15.4% of living patients (10/65) compared with 2.2% of deceased patients (1/45) (p = 0.026). *FANCA* alterations also showed a similar trend, although this did not reach statistical significance, being detected in 9.2% of living patients (6/65) and in none of the deceased patients (0/45) (p = 0.079) (**Fig. 3E**). Notably, this pattern is consistent with our *in vivo* screen, in which animals injected with *Fanca*-deficient AKPS cells also exhibited significantly prolonged survival after ICB treatment (**Suppl. Fig. 5**). No other genes in our panel showed significant differences in altered gene status between living and deceased patients, likely due in several cases to the low number of patients harboring alterations in those genes. Together, these findings support a potential association between defects in homologous recombination and Fanconi anemia pathway components and improved outcome to ICB in colorectal cancer.

### Sustained ATM inhibition induces a DNA damage–associated immune transcriptional program in MSS cells

While BRCA1 and BRCA2 mutational status may serve as prognostic biomarkers for response to immune checkpoint blockade in colorectal cancer, some other genes included in our panel represent potentially actionable therapeutic targets. Notably, ATM, a central kinase of the DNA damage response, emerged among the strongest responders to ICB in our initial *in vivo* screen (**Fig. 2B and 2D**). Given that ATM can be pharmacologically targeted with small-molecule inhibitors currently in clinical development, we sought to determine whether pharmacologic ATM inhibition could recapitulate this phenotype. To this end, we investigated the effects of two ATM inhibitors in colorectal cancer cells: AZD0156(33), the first selective ATM inhibitor to enter clinical evaluation, and AZD1390, a next-generation ATM inhibitor optimized for blood–brain barrier penetration(34). Our overarching hypothesis was that therapeutic inhibition of ATM could enhance tumor sensitivity to immunotherapy.

ATM inhibition rarely compromises cell viability unless it occurs in the context of additional defects in DNA repair pathways. Consistent with this, we did not observe a marked reduction in cell proliferation in the human colorectal cancer cell lines SW837 and LS-174T (**Fig. 4A and Suppl. Fig. 8A**) at concentrations considered to be on-target, as evidenced by suppression of downstream ATM signaling in response to the topoisomerase I inhibitor camptothecin (**Fig. 4B and Suppl. Fig. 8B**).

**Figure 4.**
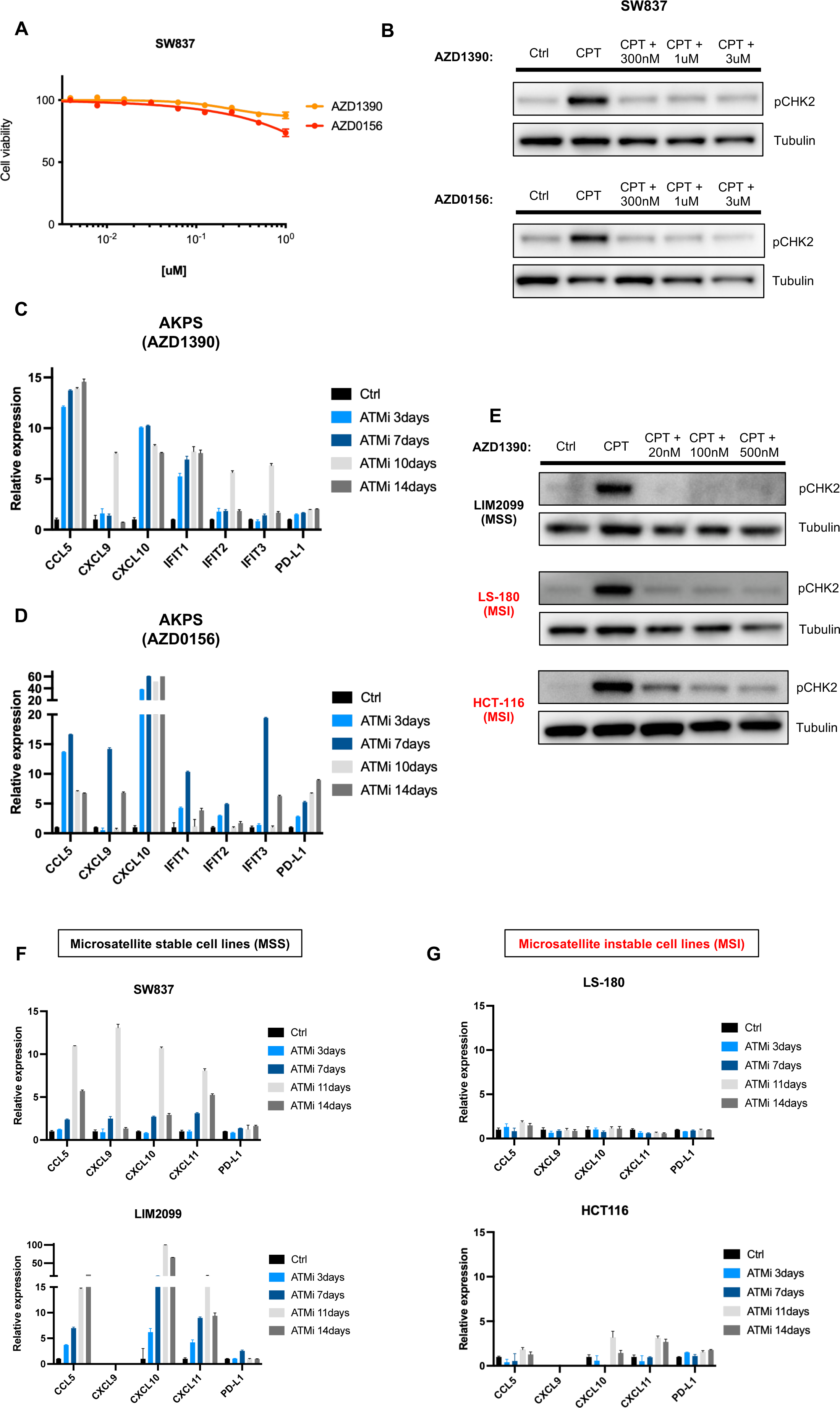
Sustained ATM inhibition induces a DNA damage–associated immune transcriptional program in MSS cells. (**A**) Cell viability of SW837 cells treated with the indicated ATM inhibitor for 72 hours. Data are normalized to vehicle-treated cells. Error bars, SD (n = 3). (**B**) Western blot analysis of SW837 cells pretreated with the indicated ATM inhibitor for 2 hours and camptothecin (CPT) for 2 hours. (**C-D**) Gene expression changes in AKPS cells treated with AZD1390 (**C**) or AZD0156 (**D**) at 500nM for the indicated times. Data are normalized to vehicle-treated cells. Error bars, SD (n = 3). (**E**) Western blot analysis of human MSS and MSI colorectal cancer cell lines pretreated with AZD1390 for 2 hours and camptothecin (CPT) for 2 hours. (**F-G**) Gene expression changes in human MSS (**F**) and MSI (**G**) colorectal cancer cell lines treated with AZD1390 (500nM) for the indicated times. Data are normalized to vehicle-treated cells. Error bars, SD (n = 3).

Despite the minimal impact of ATM inhibition on cell proliferation (**Suppl. Fig. 9**), we observed that culturing AKPS cells under ATM-inhibited conditions led to the upregulation of cytokines and interferon-stimulated genes involved in tumor–immune communication and immune cell recruitment, including CCL5, CXCL9, CXCL10, and the interferon-induced genes IFIT1, IFIT2, and IFIT3 (**Fig. 4C-D**). Notably, the transcriptional induction of these genes was detectable as early as three days after treatment in some cases, whereas in others the effect became more pronounced only after prolonged exposure, typically following 7–10 days of continuous ATM inhibition, suggesting cell adaptation under ATM inhibition selective pressure.

Although they represent only a minority of colorectal cancer cases, tumors with microsatellite instability (MSI) have been shown to be highly sensitive to immune checkpoint blockade (ICB). The underlying mismatch repair deficiency leads to persistent DNA damage and elevated mutational burden, resulting in the accumulation of neoantigens that enhance tumor immunogenicity. In addition, genomic instability in these tumors can activate innate immune sensing pathways and viral mimicry programs, promoting interferon signaling and recruitment of immune cells, both of which contribute to immune-mediated tumor elimination.

Given the transcriptional and immunomodulatory effects observed in MSS AKPS cells following prolonged ATM inhibition, we hypothesized that disruption of ATM-mediated DNA damage signaling could partially recapitulate immune-relevant features of mismatch repair–deficient tumors. Specifically, impaired ATM signaling may increase unresolved DNA damage and activate innate immune pathways, thereby promoting a more inflammatory tumor-cell state that could enhance sensitivity to anti–PD-1 therapy.

To test this hypothesis, we further evaluated transcriptional changes in both MSS and MSI human colorectal cancer cell lines following persistent pharmacological inhibition of ATM. In both groups, cells retained the ability to respond to exogenous DNA damage, as evidenced by increased phosphorylation of CHK2, a downstream target of ATM (**Fig. 4E**). However, the transcriptional response differed markedly between MSS and MSI models. In MSS human CRC cell lines (**Fig. 4F**), ATM inhibition led to a clear upregulation of cytokines involved in immune cell recruitment, including CCL5, CXCL9, CXCL10, and CXCL11. In contrast, MSI colorectal cancer cell lines exhibited only minimal transcriptional changes in response to ATM inhibition (**Fig. 4G**). These findings could suggest that the immunomodulatory effects of ATM inhibition are more evident in models with lower baseline immune activation. Collectively, our results support the notion that persistent ATM inhibition may induce selected immune-related transcriptional programs in MSS tumors that overlap with those observed in MSI tumors, thereby potentially enhancing responsiveness to anti–PD-1 blockade.

### ATM inhibition pre-treatment sensitizes tumors to PD-1 blockade *in vivo*

The ATM inhibitor AZD1390 has demonstrated promising preclinical activity, particularly in combination with radiotherapy, and is currently undergoing clinical evaluation. To test our hypothesis that ATM inhibition could partially reprogram microsatellite-stable (MSS) tumors toward a more immune-responsive transcriptional state, we designed an *in vivo* experiment in which tumor-bearing animals were treated with AZD1390 for seven days prior to initiating combination therapy with anti–PD-1 antibodies (**Fig. 5A**). This pre-treatment phase was intended to allow sustained inhibition of ATM signaling, which is expected to promote the accumulation of DNA damage and activate immune-related transcriptional programs, in accordance with our *in vitro* experiments in MSS colorectal cancer cell lines.

**Figure 5.**
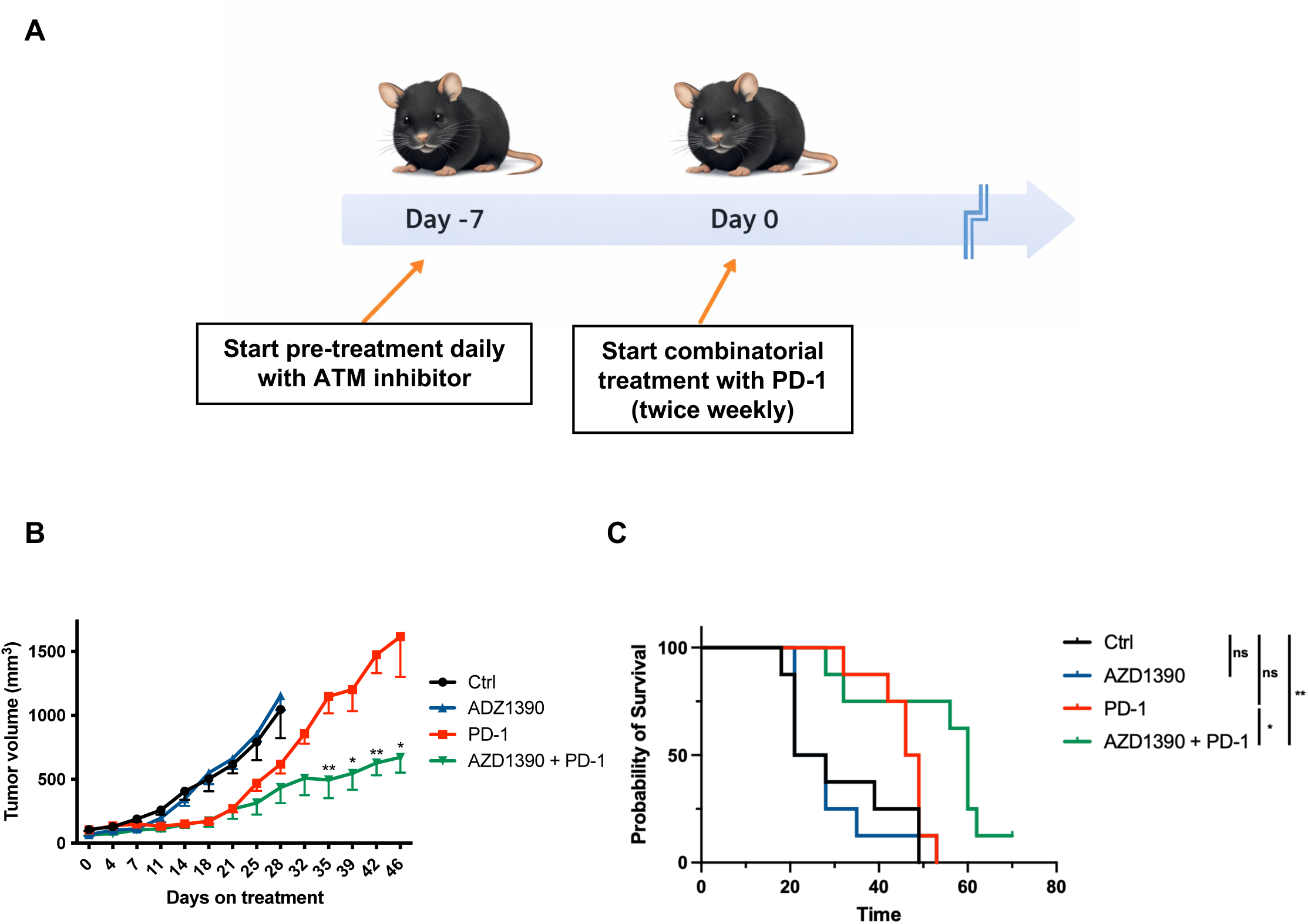
ATM inhibition pre-treatment enhances tumor sensitivity to PD-1 blockade in vivo. (**A**) Diagram depicting the experimental setup for *in vivo* experiment combining the ATM inhibitor AZD1390 with immune checkpoint blockade. (**B**) AKPS xenografts treated with AZD1390 (daily), anti-PD-1 antibodies (twice per week) or both; statistical significance was evaluated by Mann–Whitney test where *P < 0.05, **P < 0.01. (**C**) Kaplan-Meier survival curve of mice injected with AKPS cells and treated with AZD1390 (daily) or anti-PD-1 antibodies (twice per week) (n=8). *P<0.05, **P<0.01, Log rank Mantel-Cox test.

Consistent with our *in vitro* observations, treatment with AZD1390 as monotherapy did not produce measurable inhibitory effects on tumor growth *in vivo*. Treatment with anti–PD-1 antibodies initially resulted in partial tumor growth control; however, this effect was not sustained, and tumors resumed progression after approximately 21 days. In contrast, animals that received a seven-day pre-treatment with AZD1390 prior to the initiation of anti–PD-1 therapy exhibited a significant delay in tumor growth (**Fig. 5B and Suppl. Fig. 10**). This effect was accompanied by a corresponding improvement in overall survival (**Fig. 5C**). Although the combination regimen did not lead to complete tumor regressions, the magnitude of the observed benefit is notable given the absence of antitumor activity with ATM inhibition alone in this model. Collectively, these findings indicate that disruption of the ATM–CHK2 signaling axis could modulate the sensitivity of colorectal tumors to anti–PD-1 blockade *in vivo*.

## Discussion

In this study, we established AKPS as a genetically tractable and immunocompetent murine model of colorectal cancer suitable for studying tumor-intrinsic determinants of response to immune checkpoint blockade. In contrast to commonly used syngeneic colorectal cancer models, AKPS combines defined oncogenic alterations with compatibility for genome editing, enabling systematic interrogation of how specific genetic perturbations influence tumor–immune interactions *in vivo*. Using this platform, we performed an exploratory assessment of DNA damage response (DDR) pathways in colorectal cancer and found that distinct DDR defects were associated with divergent responses to PD-1 blockade, nominating selected repair alterations as candidates for biomarker development or therapeutic combination strategies.

Although defects in multiple DDR genes would be expected to increase the burden of unrepaired DNA lesions over time, responses to PD-1 blockade were not uniform across the panel. Instead, disruption of distinct repair pathways produced markedly different outcomes, ranging from enhanced sensitivity to overt resistance. This divergence suggests that the immunologic consequences of DNA repair deficiency cannot be inferred solely from the accumulation of DNA damage. Rather, the nature of the underlying lesion, the timing at which it arises, and the cellular programs engaged in response to that lesion appear to shape whether a tumor evolves toward an immune-permissive or immune-refractory state. Conceptually, these findings could support a more nuanced view in which DDR defects do not form a single biologic category in the context of immunotherapy but instead represent mechanistically distinct states with different consequences for tumor–immune interactions.

This interpretation is consistent with, but could also extend beyond, the current clinical framework for immunotherapy response in colorectal cancer. Mismatch repair deficiency and MSI-H status remain the most established genomic predictors of response to PD-1 blockade in colorectal cancer. In the phase III KEYNOTE-177 trial(19), pembrolizumab significantly improved progression-free survival compared with chemotherapy as first-line treatment for MSI-H/dMMR metastatic colorectal cancer, reinforcing dMMR/MSI-H as the clearest clinically validated immunotherapy-sensitive state in this disease. Conversely, most MSS/pMMR colorectal cancers exhibit limited sensitivity to PD-1 monotherapy, highlighting the need to define additional tumor-intrinsic mechanisms capable of generating immune responsiveness. In this context, the response patterns observed in AKPS were not explained by an MMR-like phenotype alone. Rather, integration of WES-derived genomic features with *in vivo* treatment response suggested that anti–PD-1 sensitivity was enriched among BRCA1-, ATM-, and BRCA2-deficient tumors, which also ranked among the highest for protein-altering coding mutational burden. In contrast, ATR-, MSH6-, and CDK12-deficient tumors showed poor response despite disruption of genes involved in genome maintenance. These findings indicate that the presence of a DDR defect may not be sufficient to predict immunotherapy sensitivity and that the immunologic consequences of repair deficiency could depend on the specific pathway perturbed and the type of genomic state that emerges during tumor evolution.

The overall survival data in our *in vivo* study further supported a pathway-specific model. The greatest survival benefit was observed in FANCA-, BRCA1-, and BRCA2-deficient tumors, implicating the HR/Fanconi axis. Notably, FANCA did not show high protein-altering coding mutational burden but instead displayed prominent CNV burden and loss-dominated copy-number architecture. This may suggest that large-scale genomic instability may contribute to therapeutic benefit independently of SNV-derived coding mutation burden. Such a model is biologically plausible because chromosomal instability can generate micronuclei and cytosolic DNA, thereby activating cGAS–STING and other innate immune pathways(35), although the consequences of chronic chromosomal instability can be context dependent and may also support tumor adaptation. Together, these observations suggest that anti–PD-1 response in AKPS cannot be explained by total mutational burden alone. Instead, therapeutic sensitivity may reflect distinct consequences of DDR disruption, including coding mutational burden, copy-number alterations, and activation of innate immune programs.

The strong phenotype observed with BRCA1 and BRCA2 loss is particularly informative in this regard. Both genes are core components of homologous recombination repair, and their disruption was associated with the most robust responses to PD-1 blockade in our screen, including frequent tumor regressions and prolonged survival. By contrast, perturbation of other HR-associated genes, such as PALB2, was associated with a more modest phenotype. One interpretation is that disruption of the central BRCA1/BRCA2 axis produces a more profound and durable impairment of homologous recombination than loss of accessory regulators. This may lead to the accumulation of DNA lesions or repair-associated genomic alterations that are more effective at promoting immune recognition. The observation that BRCA1- and BRCA2-deficient tumors also displayed increased expression of chemokines involved in immune cell recruitment supports this idea and suggests that not all HR-related defects are equivalent in their capacity to generate an inflamed tumor state. More broadly, these data raise the possibility that the therapeutic relevance of HR deficiency in colorectal cancer may depend on where the perturbation lies within the pathway.

The differences observed between BRCA1 and BRCA2 across experiments are also worth considering. In the initial screen, BRCA1 deficiency appeared to confer the stronger phenotype, whereas in the validation cohort an enhanced effect was more evident with BRCA2 loss. These differences may reflect technical factors, including sgRNA-dependent variation in editing efficiency, allelic disruption, or clonal selection during *in vitro* expansion. At the same time, they are biologically plausible in light of the nonredundant functions of BRCA1 and BRCA2 within homologous recombination. BRCA1 acts upstream in end resection and repair pathway choice, whereas BRCA2 functions more directly in RAD51 loading and strand invasion. These differences could influence not only the magnitude of repair impairment, but also its durability and the spectrum of secondary lesions that accumulate during tumor evolution. Accordingly, the immune consequences of BRCA1 versus BRCA2 loss may overlap without being identical.

The identification of ATM as a sensitizing genotype has additional translational importance because, unlike BRCA1 or BRCA2, ATM is directly druggable. In colorectal cancer cell lines, sustained ATM inhibition did not measurably impair proliferation under the conditions tested, but it did induce cytokine and interferon-related transcriptional programs, particularly in MSS models. This dissociation between growth inhibition and immunomodulatory activity is noteworthy because it suggests that the value of ATM targeting in this setting may not lie primarily in direct cytotoxicity, but in its ability to alter tumor cell state and thereby modify susceptibility to immune attack. The subsequent *in vivo* finding that ATM inhibitor pre-treatment improved the activity of anti–PD-1 therapy, despite minimal activity of the inhibitor alone, is consistent with that interpretation. Taken together, these data could support a model in which sustained ATM inhibition can function as an immune-priming intervention in MSS colorectal cancer.

An important conceptual implication of these results is that induction of a more immunogenic state in MSS colorectal cancer may be achievable without fully reproducing an MSI tumor. In our hands, ATM inhibition did not simply convert MSS cells into MSI equivalents; rather, it induced selected transcriptional features associated with immune activation, including expression of chemokines involved in leukocyte recruitment. The goal of DDR-targeted immunomodulation may not be to phenocopy mismatch repair deficiency in a strict molecular sense, but to induce enough of its immune-relevant features to lower the threshold for response to checkpoint blockade. That effect may be particularly valuable in MSS/pMMR colorectal cancers, where baseline sensitivity to PD-1 monotherapy is limited. From a therapeutic perspective, these data therefore support investigation of ATM inhibition not only as a radiosensitizing or DNA damage–enhancing strategy, but also as a means of reshaping the tumor microenvironment in otherwise poorly responsive disease.

The poor response of ATR-deficient tumors in our screen is also notable because it illustrates that disruption of a DDR kinase does not necessarily phenocopy disruption of a related pathway component. This is especially interesting given the established crosstalk between ATM and ATR signaling and prior reports suggesting that ATR inhibition can enhance checkpoint immunotherapy in other settings. Chronic genetic ATR loss in our model may select for tumor cell populations with adaptive changes distinct from those induced by transient pharmacologic inhibition. Alternatively, ATR deficiency may generate a pattern of replication stress and downstream signaling that favors immune evasion or reduced antigenicity in this specific colorectal cancer context. More generally, these findings emphasize that conclusions drawn from pharmacologic inhibition of a DDR kinase in one tumor type cannot necessarily be extrapolated to constitutive genetic disruption in another. The immune consequences of DDR perturbation appear to be highly dependent on context, duration, and evolutionary selection.

The analysis of the MSK-IMPACT cohort provides preliminary clinical support for our experimental observations. Within the genes represented in our panel, BRCA1 alterations were significantly associated with survival status, and FANCA alterations showed a similar trend. The concordance of the FANCA signal with our *in vivo* findings is encouraging, particularly because FANCA occupies a related biologic space at the intersection of the Fanconi anemia and homologous recombination pathways. However, the number of altered cases was limited, and the available clinical annotation was restricted to living versus deceased status rather than detailed time-to-event analysis. These data therefore do not establish BRCA1 or FANCA as validated predictive biomarkers, but they do strengthen the rationale for further clinical evaluation of selected DDR alterations as candidate modifiers of immunotherapy response in colorectal cancer.

Several limitations of the study should also be considered. First, although the AKPS model allows controlled genetic perturbation, the edited populations were expanded for several weeks before implantation in order to permit divergence in repair-deficient trajectories. This design was intentional and biologically relevant, but it also means that each genotype likely represents a heterogeneous population of subclones shaped by both the targeted gene disruption and subsequent selection. As a result, some phenotypes may reflect emergent properties of evolution under a given repair defect rather than the immediate consequence of gene loss alone. Second, the study focuses primarily on tumor-intrinsic transcriptional changes and treatment outcomes, leaving the precise immune cell populations responsible for the observed responses to be defined in greater detail.

Overall, our findings support a framework in which the effect of DNA repair deficiency on checkpoint blockade response may be pathway specific. In colorectal cancer, this distinction is likely to be clinically relevant both for biomarker development and for therapeutic design. The data suggest that selected homologous recombination and Fanconi pathway defects, particularly those involving the BRCA axis, may identify tumors with enhanced immune sensitivity, whereas pharmacologic targeting of ATM may provide a means to induce a more permissive immune state in otherwise refractory MSS disease. More broadly, this work suggests that rational combinations of DDR-targeting agents with immune checkpoint blockade should be guided by the specific repair pathway altered and the biological consequences of that alteration, rather than by the assumption that all forms of DNA repair deficiency or mutation accumulation converge on the same immunologic phenotype.

