## Supplementary Figures for "Genetic dissection of DNA damage response pathways reveals divergent modulation of PD-1 blockade response in colorectal cancer"

#### Supplementary Figure S1

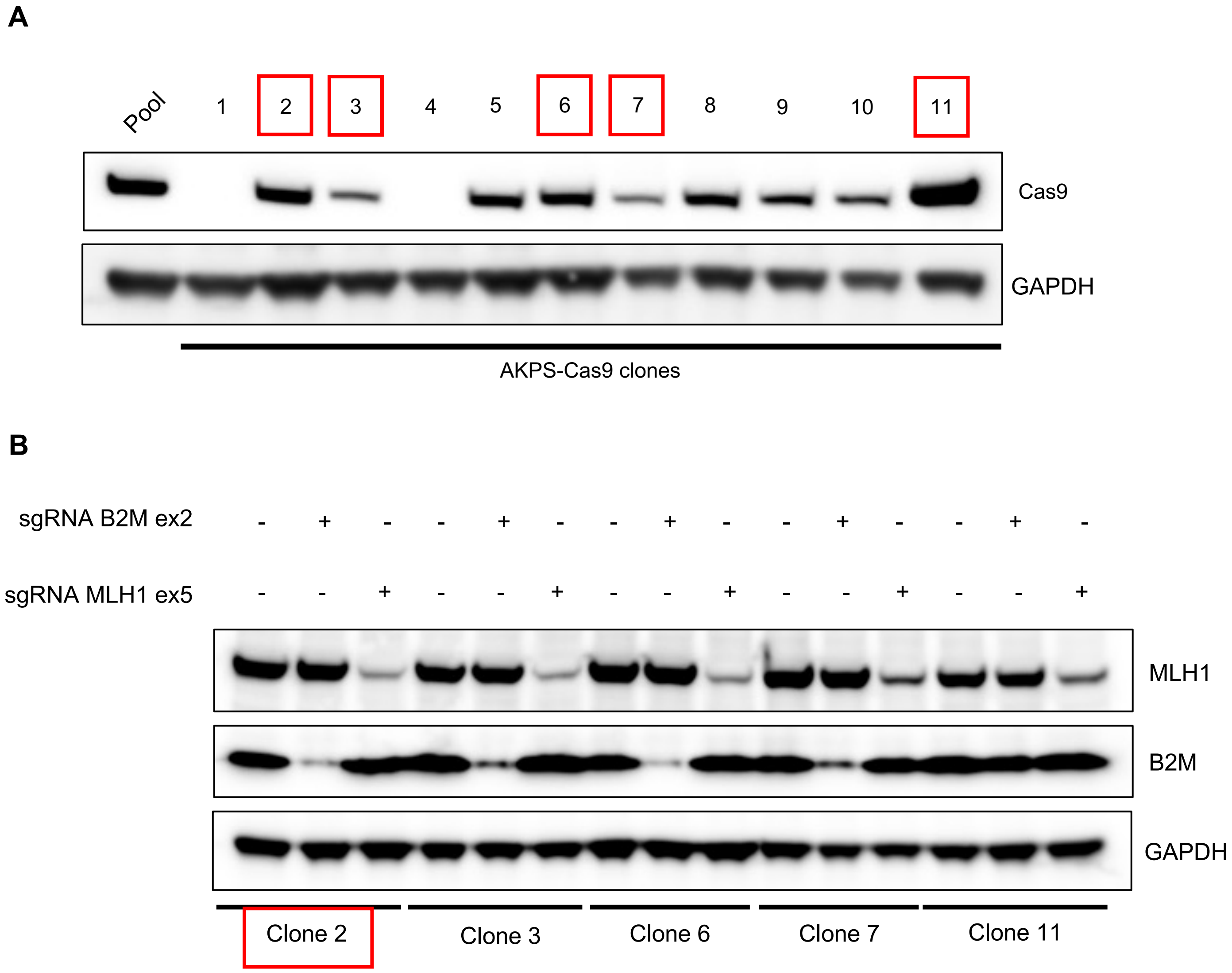


**Supplementary Figure S1 | Isolation and functional validation of Cas9-expressing AKPS clones. A**, Western blot analysis of Cas9-expressing AKPS clones. **B**, Western blot analysis of Cas9-expressing AKPS clones transduced with the indicated sgRNA.

#### Supplementary Figure S2

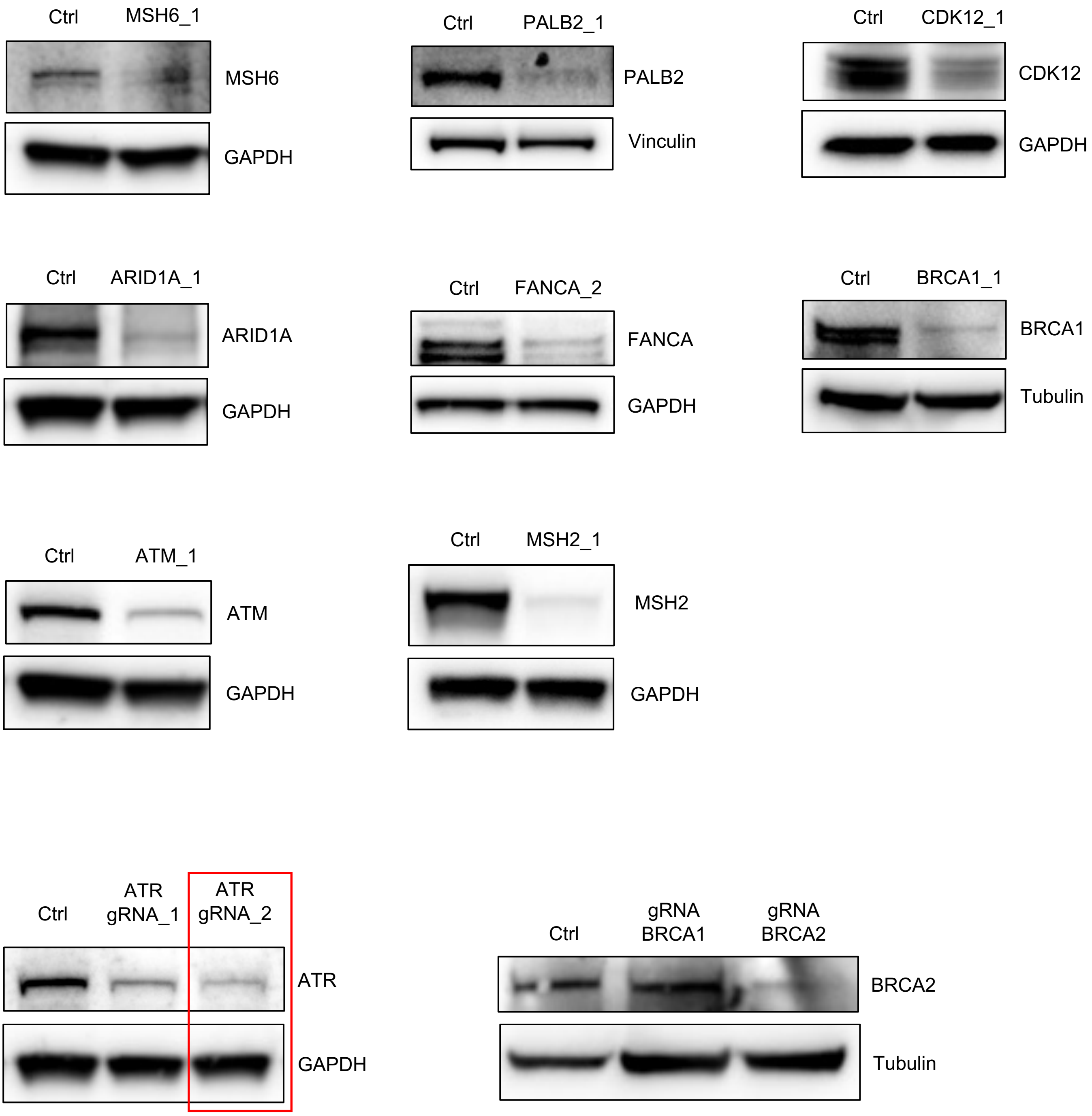


**Supplementary Figure S2 | Functional validation of vectors used for isogenic cell line panel.** Western blot analysis of AKPS-Cas9 clone 2 cells infected with indicated gRNA vector. For ATR, gRNA_2 was used in the *in vivo* screen. Since the antibody for mouse BRCA2 is not commercially available, cells expressing a BRCA1-targeting gRNA were included alongside non-targeting control cells as an additional specificity control for identification of the correct BRCA2 band.

#### Supplementary Figure S3

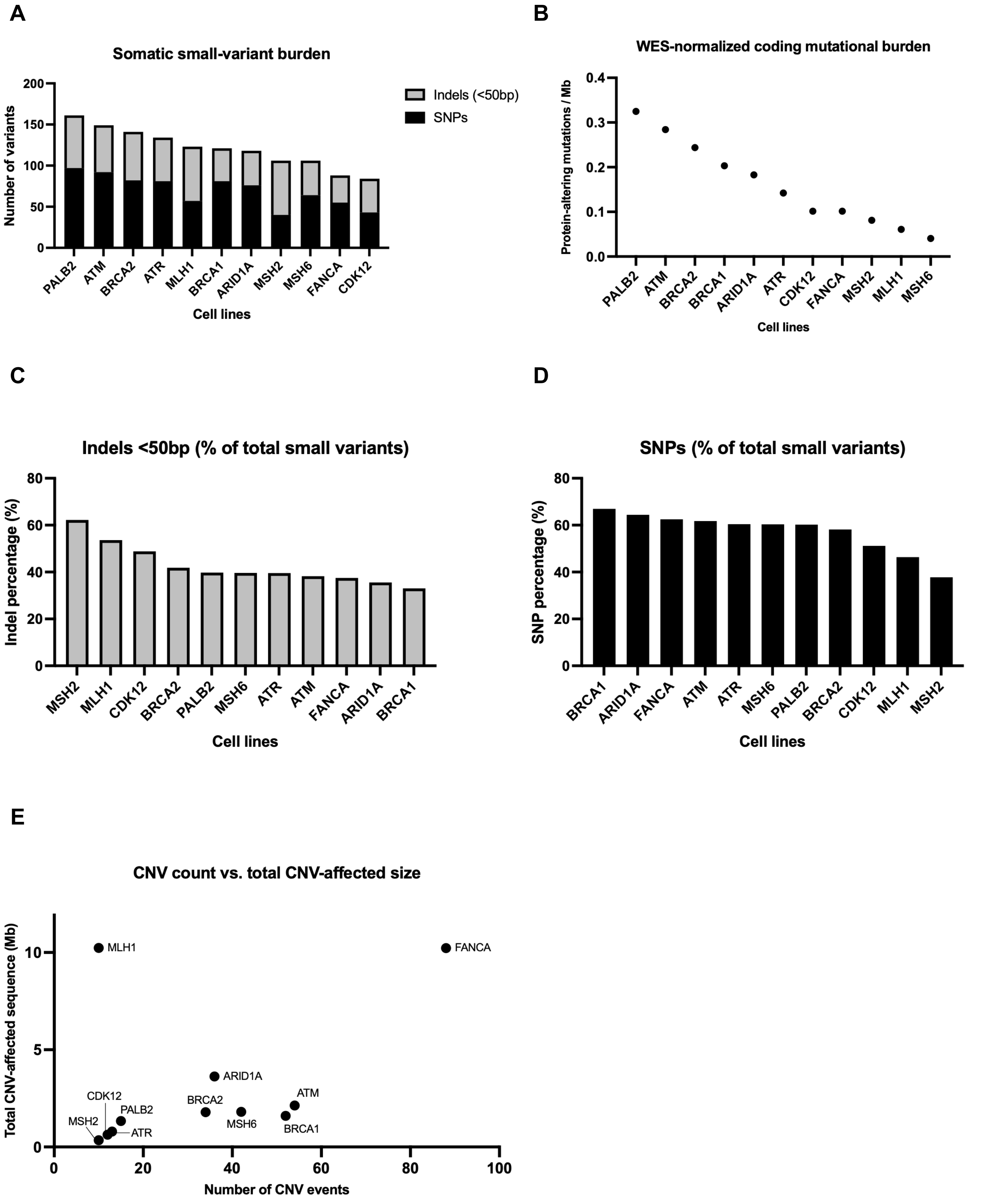


**Supplementary Figure S3 | Genomic alteration profiles of AKPS isogenic knockout cell populations after *in vitro* evolution.** Somatic genomic alterations were assessed by whole-exome sequencing (WES) in isogenic knockout cell populations expanded *in vitro* for 4–5 weeks following CRISPR-mediated perturbation. The parental control cell line was used as the matched reference for somatic variant calling. **A,** Somatic small-variant burden per knockout population, shown as the number of somatic SNPs detected by MuTect and somatic indels detected by Strelka. Bars are ranked by total small-variant burden. **B**, WES-normalized coding mutational burden, calculated as the number of protein-altering coding somatic small variants divided by the sample-specific covered WES target territory in megabases. Protein-altering coding variants included nonsynonymous SNVs, stop gain SNVs, stoploss SNVs, and coding indels. CNVs were not included in the coding mutational burden calculation. **C**, Indels expressed as a percentage of total somatic small variants, calculated as indels/(SNPs + indels) × 100. **D**, SNPs expressed as a percentage of total somatic small variants, calculated as SNPs/(SNPs + indels) × 100. **E,** Relationship between total CNV event count and total CNV-affected genomic territory for each knockout population. Each point represents one knockout population.

Supplementary Figure S4

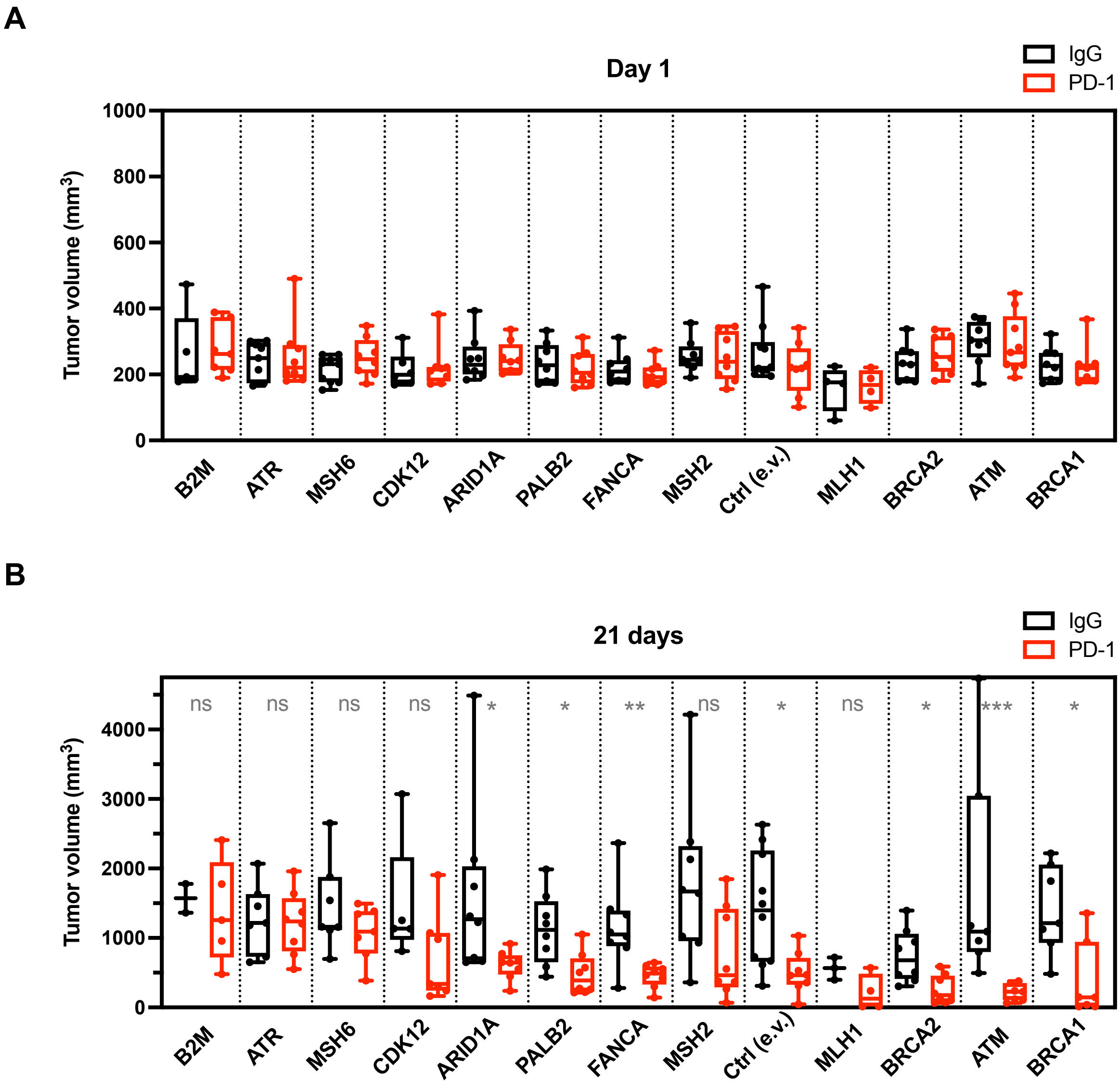


**Supplementary Figure S4 | Alterations in DDR genes confer different degree of response to PD-1 in colorectal cancer.** Shown is tumor volumes for our panel of AKPS isogenic cell lines on the first day of treatment (**A**) and on day 21 (**B**). *, $p<0.05$; **, $p<0.01$; ***, $p<0.001$ (Mann-Whitney test)

#### Supplementary Figure S5

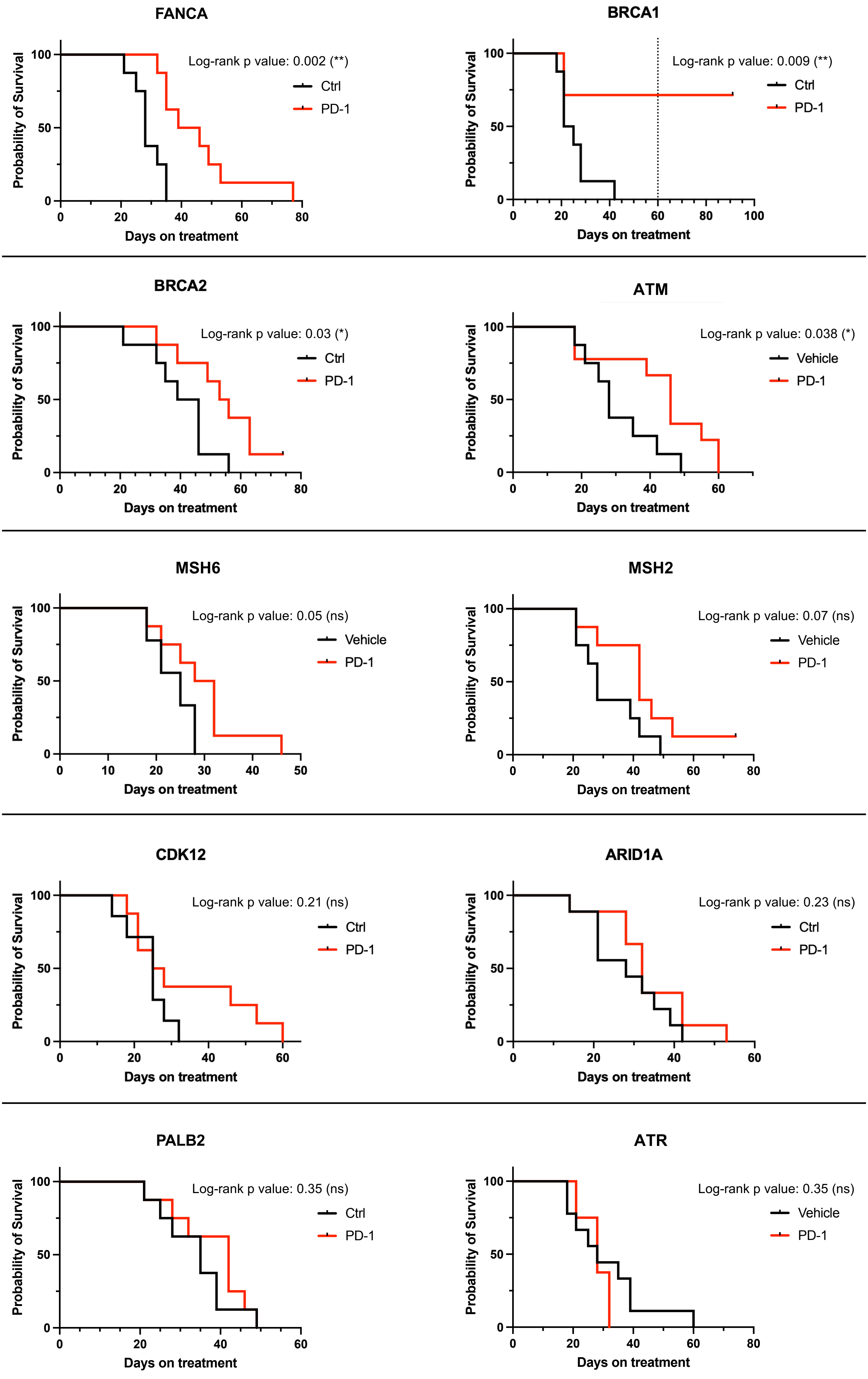


**Supplementary Figure S5 | Alterations in DDR genes confer different survival advantage in response to PD-1 blockade in colorectal cancer.** Kaplan-Meier survival curves of mice injected with AKPS cells expressing gRNAs targeting the indicated genes and treated with anti-PD-1 antibodies (10 mg $kg^{-1}$ twice per week) or an IgG control. Depicted is Log rank Mantel-Cox test.

#### Supplementary Figure S6

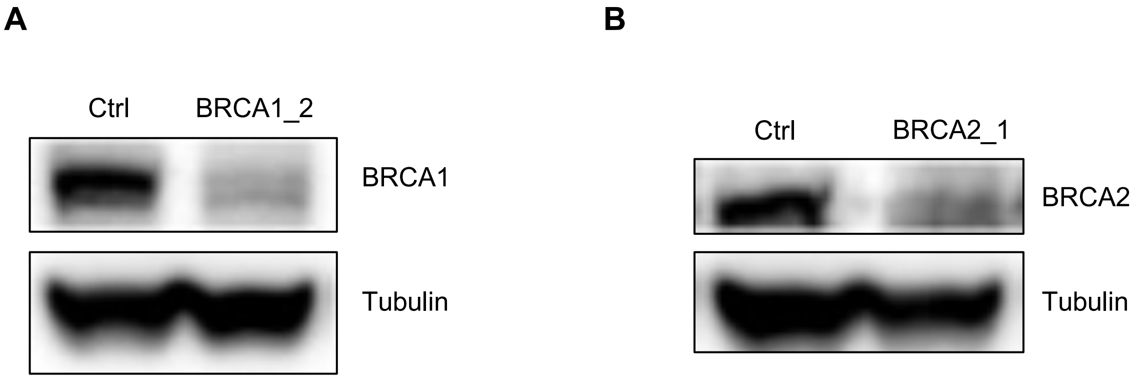


**Supplementary Figure S6 | Characterization of vectors used to independently validate BRCA1- and BRCA2-loss of function phenotype.** Western blot analysis of AKPS-Cas9 clone 2 cells infected with a gRNA vector targeting BRCA1 (**A**) or BRCA2 (**B**).

#### Supplementary Figure S7

**A**

**Colorectal (n=110)**

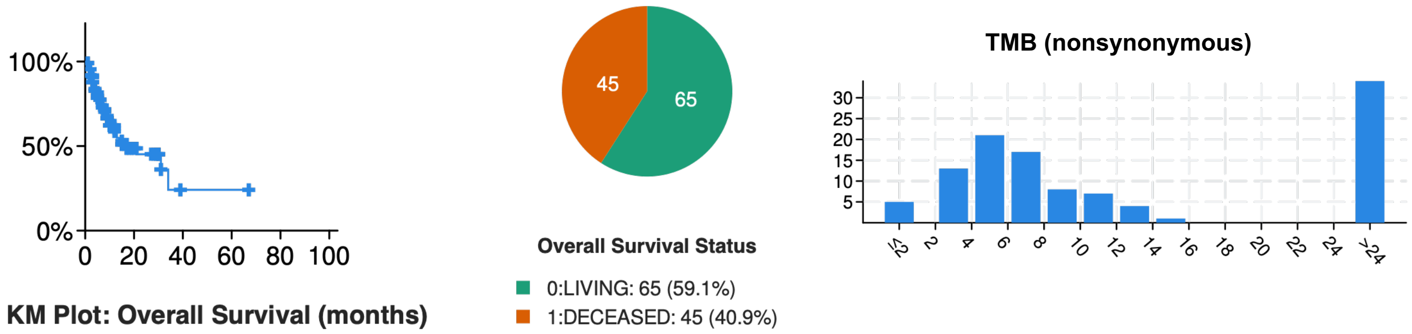


**B**

**Only BRCA1-altered colorectal (n=10)**

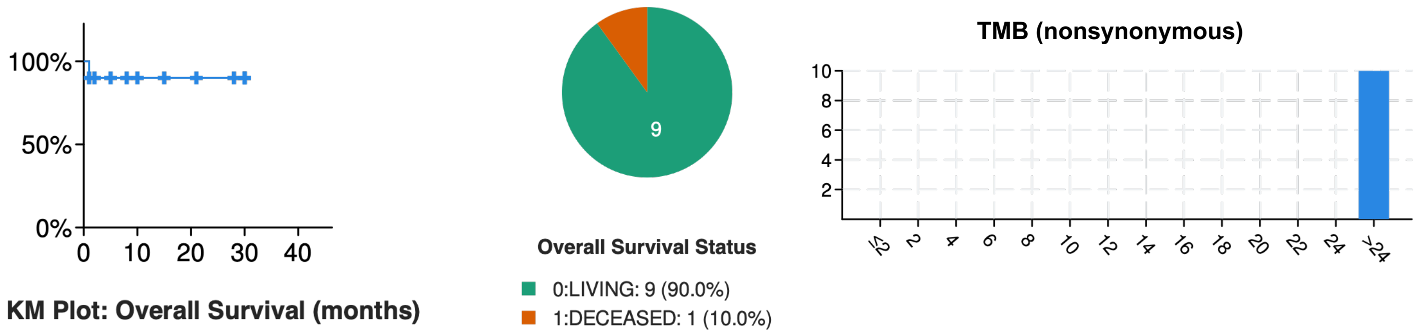


**Supplementary Figure S7 | Analysis of the colorectal cancer patients treated with immune checkpoint blockade in the MSK-IMPACT database.** Data displayed includes survival analysis and tumor mutational burden for the entire colorectal cancer cohort (**A**) or specifically the BRCA1-altered patients (**B**).

#### Supplementary Figure S8

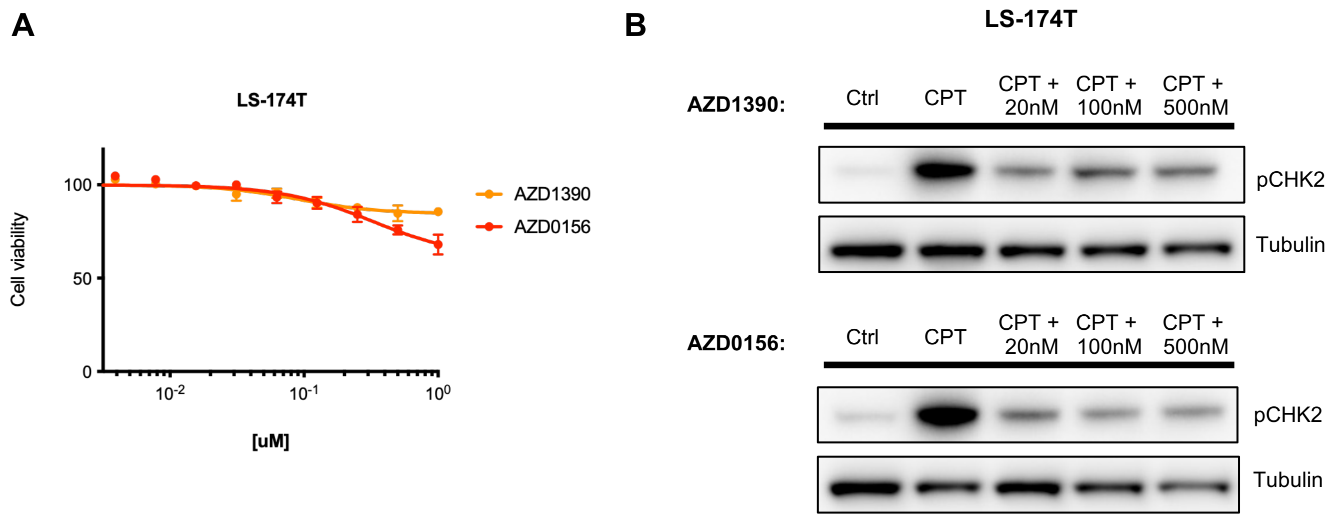


**Supplementary Figure S8 | ATM inhibition does not significantly compromise cell viability of the MSI-positive cell line LS-174T. A**, Cell viability of LS-174T cells treated with the indicated ATM inhibitor for 72 hours. Data are normalized to vehicle-treated cells. Error bars, SD (n = 3). (**B**) Western blot analysis of LS-174T cells pretreated with the indicated ATM inhibitor for 2 hours and camptothecin (CPT) for 2 hours.

#### Supplementary Figure S9

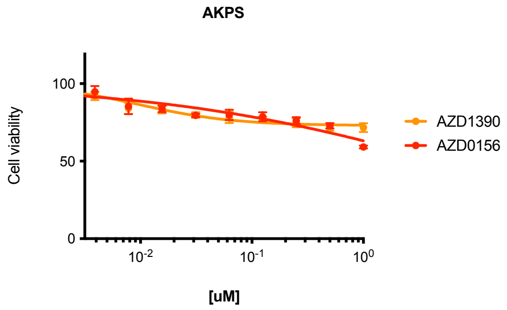


**Supplementary Figure S9 | ATM inhibition does not significantly compromise cell viability of mouse colorectal cancer cells.** Cell viability of AKPS cells treated with the indicated ATM inhibitor for 72 hours. Data are normalized to vehicle-treated cells. Error bars, SD (n = 3).

### Supplementary Figure S10

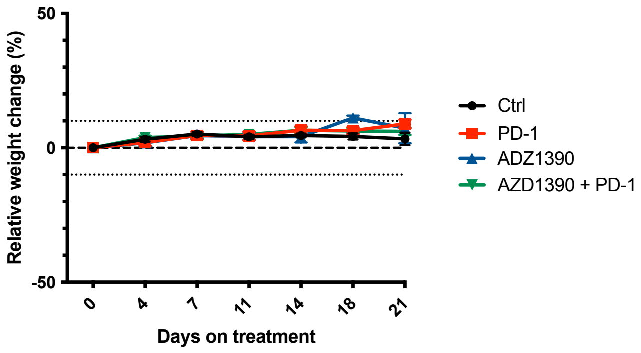


**Supplementary Figure S10 | Percent body weight change for xenograft models from Figure 5B.**
