## Supplementary Tables for "Genetic dissection of DNA damage response pathways reveals divergent modulation of PD-1 blockade response in colorectal cancer"

**Supplementary Table 1**

| Gene | Sequence |
|---|---|
| ATM | GGATCACGGAGTACATCCAGGGG |
| ATR | TGAAGATGGATTCGTAAAAGAGG |
| ARID1A | CAGCAGAACTCGCACGACCACGG |
| B2M | AGTATACTCACGCCACCCACCGG |
| BRCA1_gRNA1 | GCTGATACAATGTGTGATAGAGG |
| BRCA1_gRNA2 | GAGCTTCAAATCGATAGTTGTGG |
| BRCA2_gRNA1 | AAATGAGAATGGGCAAGACAAGG |
| BRCA2_gRNA2 | TGCTTTGGCATATTATACGGAGG |
| CDK12 | GTCCTTACAGCCGACGACAGAGG |
| FANCA | TGTGTTAAGATTGTTCCCAGAGG |
| MLH1 | CAACCAGGGCACCCTGATCACGG |
| MSH2 | GACAAAGATTGGTTAACCAGTGG |
| MSH6 | GAGGCAAAGGATCTCAACGGAGG |
| PALB2 | AGGAAATACTGTCTCCGCGGAGG |

**Supplementary Table 1 | List of genes targeted and the CRISPR sgRNA sequences**

**Supplementary Table 2**

| Patient | Follow-up time | BRCA1 status | MLH1 status | POLE status | TMB |
|---|---|---|---|---|---|
| P-0001808 | 10 months (living) | K520N | - | P286R (mutator phenotype) | 178.44 |
| P-0004051 | 28 months (living) | K654Sfs*47 (del) | - | - | 68.87 |
| P-0004783 | 21 months (living) | I171T | - | - | 47.22 |
| P-0005151 | 30 months (living) | A1615V L1764* (fs del) | - | - | 65.91 |
| P-0009830 | 30 months (living) | K654Sfs*47 (del) | T117M | - | 44.27 |
| P-0017986 | 2 months (living) | R1203Q | - | - | 61.44 |
| P-0019428 | 5 months (living) | N567Ifs*5 (FS del) | - | - | 46.52 |
| P-0019475 | 1 month (deceased) | K654Sfs*47 (del) | R487* | C1357Y | 93.04 |
| P-001994 | 8 months (living) | F1036Lfs*12 (del) | - | - | 38.62 |
| P-0020399 | 15 months (living) | V340Gfs*6 (ins) | X32_splice | R616C | 61.44 |

**Supplementary Table 2 | Clinical and genomic characteristics for the BRCA1-altered colorectal cancer cohort treated with immune checkpoint blockade in the MSK-IMPACT database.**
